# FourierMIL: Fourier filtering-based multiple instance learning for whole slide image analysis

**DOI:** 10.1101/2024.08.01.606256

**Authors:** Yi Zheng, Harsh Sharma, Margrit Betke, Jennifer E. Beane, Vijaya B. Kolachalama

## Abstract

Recent advancements in computer vision, driven by convolutional neural network, multilayer perceptron and transformer architectures, have significantly improved the analysis on natural images. Despite their potential, the application of these architectures in digital pathology, specifically for analyzing gigapixel-resolution whole-slide images (WSIs), remains challenging due to the extensive and variable sizes of these images. Here we present a multiple instance learning framework that leverages the discrete Fourier transform and learns from WSIs. Dubbed as FourierMIL, our framework is designed to capture both global and local dependencies within WSIs. To validate the efficacy of our model, we conducted extensive experiments on a prevalent computational pathology challenge: tumor classification. Our results demonstrate that FourierMIL outperforms existing state-of-the-art methods, marking a significant advancement in the field of digital pathology and highlighting the potential of attention-free architectures in managing the complexities related to WSI analysis. The code will be released for public access upon the manuscript’s acceptance.

## I. Introduction

DEEP learning has revolutionized computer vision, offering significant potential for analyzing gigapixel-resolution whole-slide images (WSIs) in computational pathology. Gigapixel WSIs present a challenge due to their size, which standard vision models like ResNet [1]—designed for smaller, fixed-size inputs struggle to process without substantial computational resources. Reducing WSI dimensions to manageable sizes, such as 224 × 224 pixels, results in the loss of critical diagnostic details. To navigate this, deep learning-based multiple instance learning (MIL) has been utilized [2]–[4], treating the diagnosis as a weakly supervised learning problem. This method segments the WSI into smaller patches for extracting features with convolutional neural networks (CNNs) and then aggregates these features for a comprehensive WSI-level representation. In essence, the field has long been dominated by CNN-based methods [1], [5], [6]. However, their approach often overlooks essential contextual and hierarchical information within pathological images, crucial for tasks such as survival analysis in cancer [2], [7]. Graph convolution networks (GCNs) have introduced the ability to create hierarchical representations of image patches, offering a nuanced way to capture spatial information by representing WSIs as graph-based structures. Yet, the challenge of determining optimal edge weights and graph topology can limit the effectiveness of GCNs in fully capturing spatial relationships. More recently, transformer-based architectures [8], [9] and pure MLP-based models [10] have made breakthroughs in visual tasks. However, their application to WSIs is hindered by the large, unstructured nature of WSI patches. Here we present a multiple instance learning framework that leverages the discrete Fourier transform and captures global and local dependencies within WSIs.

We summarize the key contributions of this paper as follows:

- We propose a Fourier filtering-based multiple instance learning (FourierMIL) framework, which facilitates efficient token mixing with large dynamic kernels to capture dependencies across a large number of instances for WSI analysis.
- We introduce token padding as a solution to mitigate spectral leakage and edge effects, which arise from the non-periodic nature of token representations and the assumption of periodicity by the discrete Fourier transform.
- We formulate an MLP-based FourierMIL model that introduces a significant improvement in WSI analysis, thus surpassing the performance of existing state-of-the-art transformer and graph-based methods.

## II. Related work

### A. Multiple instance learning

Instance-level MIL strategies for WSI analysis, exemplified by the studies of Campanella et al. [11] and Xu et al. [12], employ convolutional neural networks (CNNs) where pseudo-labels are attributed to each instance derived from the overarching bag-level label, followed by the selection of the top-k instances for subsequent aggregation. Conversely, embedding-level MIL techniques, such as those demonstrated by ABMIL [2] and additional studies [13], [14], convert each WSI patch into a uniform-length embedding, from which bag-level representations are synthesized based on these individual embeddings. Certain methodologies incorporate feature clustering processes, like ClustertoConquer [15] and the method proposed by Wang et al. [16], which determine cluster centroids from feature embeddings and use these representative embeddings for final predictions. Furthermore, attention-based MIL frameworks, including ABMIL [2] and DSMIL [14], enhance instance contribution discernment by integrating trainable parameters. Specifically, ABMIL leverages a side-branch network for deriving attention weights, whereas DSMIL utilizes cosine distance to ascertain instance similarity, thereby assigning weights.

### B. Vision models for natural images

Since the introduction of AlexNet [5], CNN-based architectures have been the primary visual backbones, with VGG network [6] demonstrating state-of-the-art performance on ImageNet [17] through its use of 3 × 3 convolution and fully connected layers. ResNet [1] further advanced this field by introducing residual connections to facilitate feature transfer across layers, mitigating the issue of vanishing gradients and enhancing performance. These successes underscore the utility of locality bias and the pyramid structure of multi-stage processing for improving performance in CNN-based vision models.

The transformer architecture, initially developed for natural language processing (NLP) by Vaswani et al. [8], has recently led to significant advancements in computer vision. Dosovitskiy et al. [9] introduced the application of transformers for vision pretraining on natural images by representing 224× 224 images as sequences of 16 ×16 flattened image patches. However, the application to WSIs presents unique challenges. Unlike natural images, WSIs can be viewed as sequences or bags of patches, with an average bag size containing about 10,000 non-background 256×256 image patches at 20X magnification, significantly increasing computational complexity. This complexity renders the use of transformers and similar self-attention architectures impractical for MIL tasks related to WSIs due to the high space complexity. Kalra et al. [18] addressed this by employing set-transformers for lung cancer subtyping with bags of 100 randomly sampled histology patches. TransMIL [19] introduced a linear complexity Nyström-based attention mechanism [20] to manage the computational demands by approximating the self-attention mechanism, enabling the exploration of long-range interactions among WSI patches.

Moreover, recent works [10], [21]–[23] have explored MLP-based models by eliminating the attention mechanism in transformers. These models apply MLP layers to feature patches in spatial dimensions to aggregate spatial context. MLP-Mixer [10] uses token- and channel-mixing to capture relationships between tokens and channels. CycleMLP [21] introduces pseudo-kernels to sample tokens from various spatial locations for mixing. AS-MLP [22] and S^2^-MLP [23] utilize token shifting in vertical and horizontal directions to achieve an axial receptive field and facilitate cross-patch communication, respectively.

Building on MLP-based architectures, models that operate in the frequency domain utilize the Fourier transform to mix tokens spatially. FNet [24] mirrors the structure of MLP-mixers, with token mixing achieved through a straightforward application of the Discrete Fourier Transform (DFT), without adjusting for data distribution variations. Meanwhile, global filter networks (GFNs) [25] and adaptive Fourier neural operators (AFNO) [26] introduce a more sophisticated approach by learning Fourier filters of fixed size and performing global convolution, with AFNO further enhancing this process by incorporating dynamic filtering for global convolution tasks.

### C. Graph-based analysis

Beyond MIL and transformer-based methods, graph convolutional networks (GCNs) and other graph-oriented techniques have been employed for WSI analysis. Certain approaches concentrate exclusively on using cell identities as graph nodes [27], [28], neglecting vital prognostic tissue elements like stroma, and are constrained to analyzing limited image areas, as critiqued in [29]. Pathomic fusion [30] constructs cell-based graphs for analyzing small regions of interest (ROIs) and utilizes spectral convolutions for processing. Graph-CNN [31] selects patches within a WSI as nodes, connecting them through edges based on similarity in feature embeddings, and employs spectral convolutions for its analysis. Patch-GCN [32] generates a WSI-graph in Euclidean space using feature embeddings from all non-background patches, executing graph convolutions similar to CNNs’ local neighborhood aggregation. Alternatively, GTP [33] simplifies the nodes in a WSI-graph to a fixed count using min-cut pooling, then employs a transformer to create WSI-level representative features.

## III. Methods

### A. Preliminaries and problem formulation

We concentrate on analyzing large-scale WSIs, which we actively partition into a grid of small, distinct, and non-overlapping patches. This process allowed us to identify and exclude patches primarily containing background elements, resulting in a curated set of *L* non-background (or tissue) patches for further analysis. These patches, henceforth referred to as tokens, are each characterized by a *D*-dimensional feature vector. We utilized pre-trained feature extraction models to analyze and describe these tokens. To achieve this, we used both Resnet-50 and CTranspath [34] in our experiments. However, it is important to note that our methodology is flexible and can accommodate various feature extraction techniques. Ideally, the feature extractor should be tailored based on the specific stain used in the WSI. For instance, domain-specific models like CTranspath, which are trained on H&E stained WSIs, are expected to perform better on those slides. Instead of working with less common WSIs, a more general architecture like Resnet-50 can be employed to extract meaningful features from each patch. Consequently, the image is encapsulated as a set of features *X* = *x*_1_, …, *x*_*L*_∈ ℝ^*L*×*D*^, forming the basis for our analysis in Fig. 1(a).

**Fig. 1:**
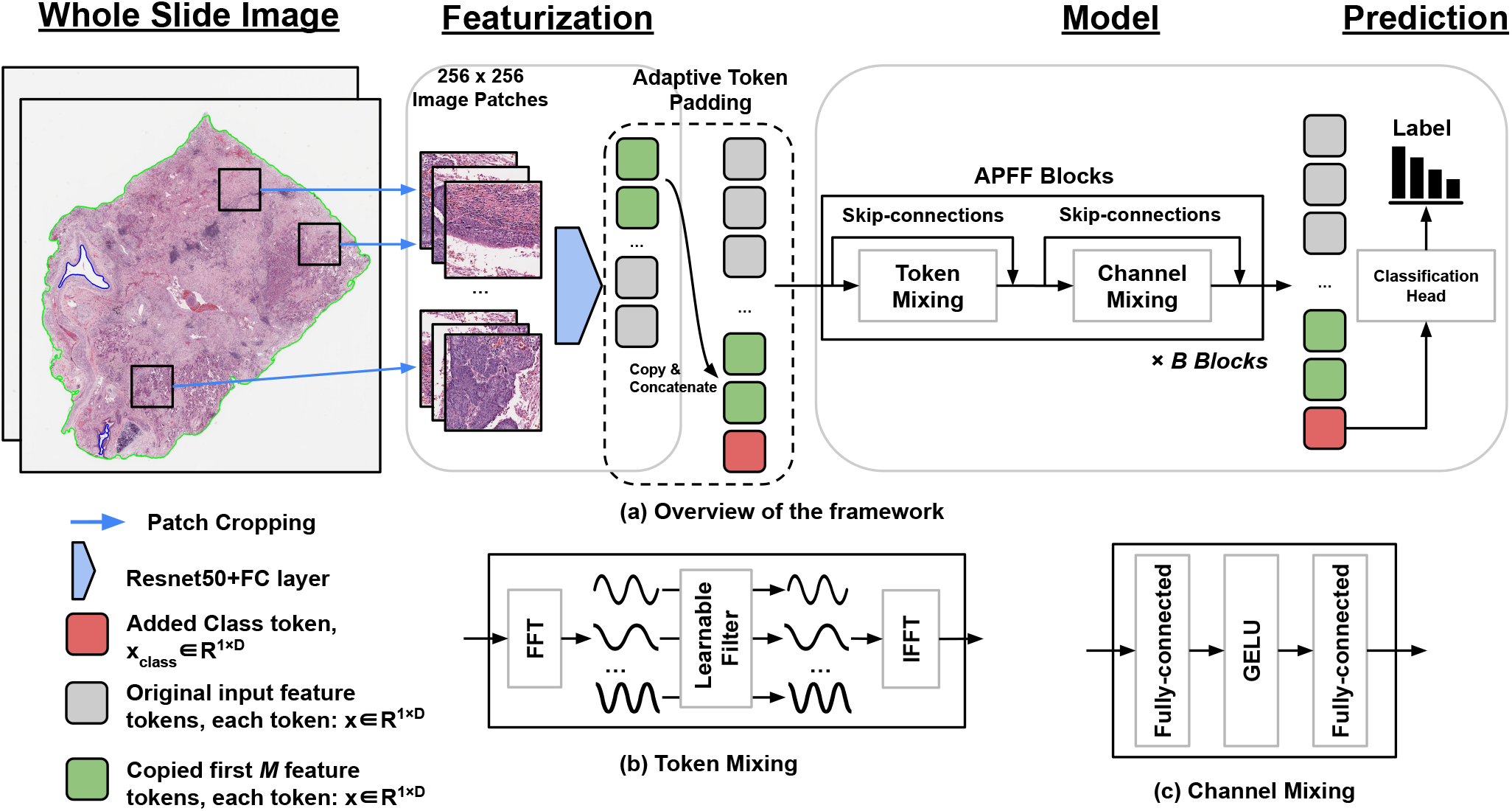
FourierMIL framework. (a) Each whole slide image is segmented into patches, excluding the background, and transformed into feature vectors via ResNet50. The patch tokens sequence is then padded, and a class token is appended, before input into the proposed model. Through a series of token mixing and channel mixing blocks, the output class token is directed to a classification head, consisting of a single linear layer, to produce a slide-level prediction. (b) Token mixing involves a fast Fourier transform (FFT), application of a learnable filter in the frequency domain, followed by an inverse fast Fourier transform (IFFT). (c) Channel mixing employs two fully-connected layers and a GELU nonlinearity, also referred to as an MLP within the paper.

Our objective is to construct a contextual embedding that not only encapsulates the intricate details within the WSIs but also demonstrates versatility and efficacy in a variety of downstream tasks. In the realm of single-label classification, our endeavor is to accurately predict the label of the entire WSI, notwithstanding the absence of individual labels for each token. This necessitates a strategic blending of the tokens, ensuring both local and global interactions are effectively leveraged to produce a comprehensive and rich representation conducive to our analytical goals.

A vision MLP-Mixer model learns to extract the rich representation of *X* with two basic operations, i.e., channel-mixing and token-mixing. The channel-mixing (CM) operation is formulated as:

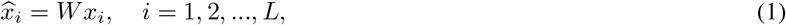

where *W* is the learnable weight and 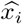 is updated token. It operates on each token independently to extract their features. To aggregate information from different tokens, the token-mixing (TM) operation can be formulated in a unified form:

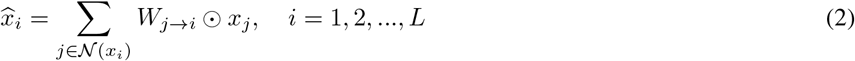

where *W*_*j*→*i*_ represents the weights of information aggregation from token *x*_*j*_ to the updated 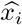, 𝒩(*x*_*i*_) is the contextual region of *x*_*i*_, and ⊙denotes Hadamard product.

The TM operation tries to capture the spatial information by mixing features from different tokens, and we revisit the prevalent TM methods in different types of network architectures in terms of their effectiveness and efficiency. For CNNs, TM is facilitated through matrix multiplication, employing deterministic network parameters as weights. Here, the kernel sizes of convolutions determine the scopes of token mixing. This makes mixing in a global scope quite costly due to the quadratically increased parameters. In transformer-like architectures, self-attention serves a similar purpose to TM in mixing tokens with pairwise correlations between query and key tokens. It however scales quadratically with the number of tokens, which impedes training high-resolution images such as WSIs. Like CNNs, MLP-Mixer-like models also mix tokens with deterministic network parameters. But they often neglect the varying semantic contents of tokens from different input images, restricting the representation ability. Our goal is then to find an alternative token-mixing strategy that achieves favorable scaling trade-offs in terms of computational complexity, memory, and downstream transfer accuracy. This requires a large 𝒩 (*x*_*i*_) and instance-adaptive *W*_*j*→*i*_ with less network parameters and low computation costs as possible.

### B. All-pass frequency filtering for token mixing

According to the convolution theorem [35], global convolution within the spatial domain equates to multiplication within the eigen transform domain. Fourier transform is an example of such eigenfunctions, so the interactions among spatial locations can be learned in the frequency domain. Based on this theorem, we note that adaptive frequency filters can serve as efficient global token mixing. In what follows, we introduce its mathematical modeling, architecture design and the equivalence between them for our proposed token mixer.

#### Modeling

We begin with an introduction to the discrete Fourier transform (DFT), a pivotal element in digital signal processing and a fundamental aspect of our framework. For clarity, considering our focus on a MIL problem where we encounter a set of tokens devoid of 2D spatial information, we concentrate on addressing these tokens in a 1D context. This approach diverges from traditional vision classification problems that typically involve input features with a 2D structure. The TM operation in Eq.(2) can be represented in a general form of global convolution, denoted as 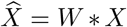. For the i-th token *X*[*i*], Eq.(2) can be reformulated in the 1D discrete form as:

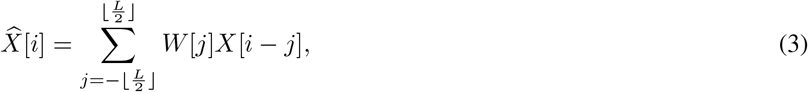

where 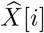 represents the updated token for *X*[*i*] after TM. *W* denotes the weights for TM, implemented by a global convolution kernel that has the same size as *X*. Based on the convolution theorem, a convolution in one domain mathematically equals the Hadamard product in its corresponding Fourier domain. Given a set of tokens, we adopt the fast Fourier transform (FFT) to obtain the corresponding frequency representations *X*_*F*_ by *X*_*F*_ = ℱ(*X*):

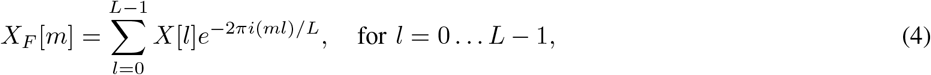

where *i* is the imaginary unit, and the features of different spatial positions in *X*_*F*_ correspond to different frequency components of *X*.

#### Token mixing

Based on the aforementioned convolution theorem, global TM for *X* is equal to filtering its frequency representation *X*_*F*_ with a learnable filter *W*_*F*_, i.e.,

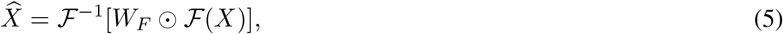

where ℱ^−1^ is the inverse FFT (iFFT), *W*_*F*_ is the Fourier transform of convolution kernel weight *W* for TM and has the same shape with *X*_*F*_.

#### Adaptive learnable filter weights

To design the filter weights *K*_*F*_, we revisit CM layer in MLP-Mixer [10]. The CM layer acts on each token of *X*, maps ℝ^*D*^ ↦ℝ^*D*^, and is shared across all tokens. It contains two fully-connected layers and a nonlinearity applied independently to each token:

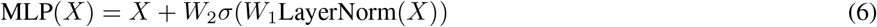

where *W*_1_ and *W*_2_ are trainable parameters, and *σ* is an element-wise nonlinearity (GELU [36]). This CM operation along the channel dimension is adopted by us as filtering in the frequency domain:

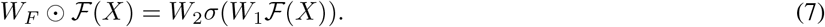

GFNet [25] learned the trainable filters *W*_*F*_ by multiplying ℱ (*X*) with fixed-shaped *W*_*F*_ through element-wise multiplication, which did not support our variable-size input tokens. Instead, the MLP-like filters *W*_1_ and *W*_2_ in eq. 7 were adaptive and invariant to any length of input tokens. To reduce the model parameters, we imposed a block diagonal structure on *W*_1_ and *W*_2_ following AFNO [26], where *W*_1_ and *W*_2_ were divided into *h* weight blocks of size *D/h*.

In existing Fourier filtering based models, such as the AFNO [26], an ideal low-pass filter is employed within the frequency domain. AFNO implemented a soft-thresholding and shrinkage operation in the frequency domain to adaptively mask tokens according to their relevance to the end task. However, the soft shrinkage operation, by applying a threshold (*λ*), suppressed small values, effectively analogous to eliminating high-frequency components while preserving low-frequency components. However, the high-frequency components of images, which encapsulate fine details, are crucial for image analysis, particularly in histopathological examinations. Therefore, in contrast to the conventional approach of utilizing low-pass or high-pass filters, our methodology involves the deployment of a filter that permits the passage of token signals across all frequencies. This token-mixing approach, which we have termed All-Pass Frequency Filtering (APFF), ensures the preservation and utilization of comprehensive detail in image analysis (see Algorithm 1).

##### Algorithm 1

All-pass frequency filtering (APFF)

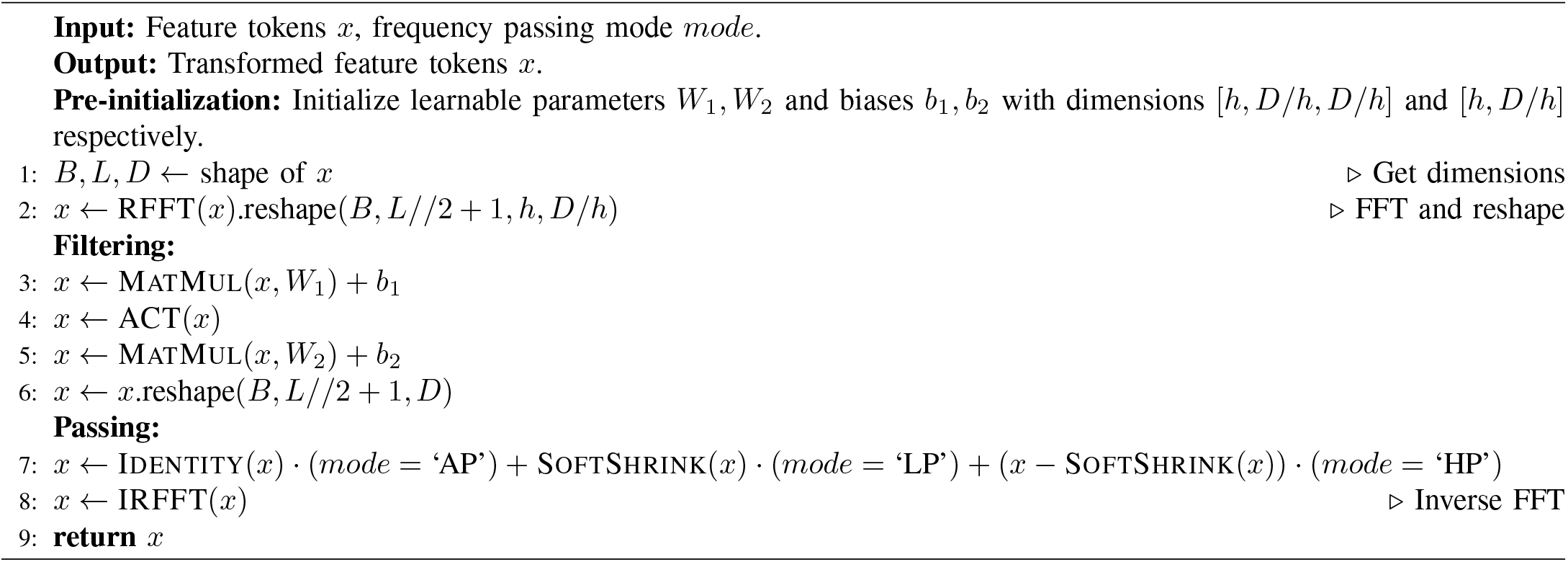

### C. Fourier filtering-based multiple instance learning

Here we introduce our MLP-Mixer-based FourierMIL model which consists of several APFF blocks. The entire architecture of a APFF block can be formulated as:

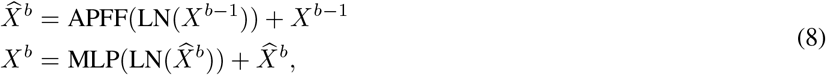

where *X*^*b*−1^ is the output of the (*b*− 1)-th APFF block. We adopted the commonly used MLP [10] in Eq. 6 and APFF with Layer Normalization (LN) for channel mixing and token-mixing, respectively. Skip-connections for channel mixing and token mixing were adopted to facilitate model training. The skip-connections for token mixing can compensate the incomplete local features caused by the Fourier transform (FT), since when the FT decomposes an image into its frequency components, local features such as edges or small objects, may not be well represented because the FT spreads the information across the entire frequency spectrum. By stacking multiple APFF blocks, we constructed our network, namely FourierMIL, as shown in Fig. 1(a).

### D. Adaptive token padding

To mitigate potential artifacts introduced by the Fourier transform without modifying the original image token’s frequency content, we introduced an Adaptive token padding (ATP) strategy applied to input *X* prior to processing with FourierMIL. ATP enhances the input for FT, leading to a more refined frequency grid in the domain. This refinement allows for a closer spacing of frequency components, facilitating more precise spectral filtering and, consequently, a more detailed convolution of tokens. Additionally, ATP acts as a form of frequency domain interpolation among original tokens, smoothing the Fourier transform of token features. This smoothing aids FourierMIL in more effectively discerning peaks and troughs within the frequency domain. Furthermore, ATP addresses the issue of spectral leakage arising from the non-periodic nature of image tokens, potentially introducing artificial high-frequency components, such as edge effects or Gibbs phenomena. These artifacts, which can obscure the true frequency content of an image, are significantly minimized, ensuring a cleaner frequency representation for enhanced image analysis. To determine the padding number of tokens, instead of using a fixed numbe√r or a fixed ratio of original tokens, we selected the padding number *M* based on the number of original tokens, i.e., 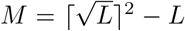, where *L* is the number of original tokens. Here *M* is a function of *L*, which is adaptive. The proposed approach can also regularize the network and improve its robustness. Additional details on ATP are provided in Algorithm 2.

#### Algorithm 2

Adaptive token padding (ATP)

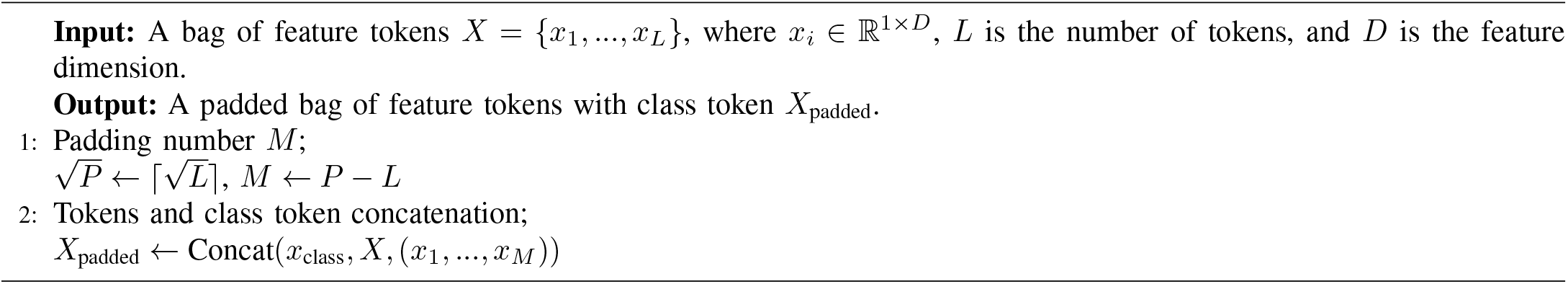

### E. Feature aggregation and slide-level prediction

Similar to ViT [9], we added a learnable class token in ATP to padded tokens, whose state at the output of FourierMIL serves as the slide-level representation. The classification head in Fig. 1(a) was implemented by a single linear layer:

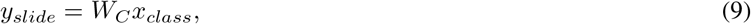

where *W*_*C*_ ∈ ℝ^*C*×*D*^ and *x*_*class*_ ∈ ℝ^1×*D*^. Here *C* corresponds to the number of classes.

## IV. Experiments

### A. Study population

We performed a series of experiments across three publicly accessible datasets: CAMELYON16 [37], The Cancer Genome Atlas (TCGA) Non-Small Cell Lung Cancer (NSCLC) [38], and the Clinical Proteomic Tumor Analysis Consortium (CPTAC) [39], as shown in Table I. For each WSI, tissue segmentation was achieved using Otsu’s method, followed by the extraction of non-overlapping patches of size 256 × 256 at 20X magnification. Feature encoding was performed using a ResNet-50 model, pre-trained on ImageNet, incorporating a 256-dimensional fully-connected layer to yield a 1024-dimensional embedding for each patch, while the ResNet-50’s parameters remained unchanged.

**TABLE I:** Study population. Whole slide images and corresponding clinical information from three distinct cohorts including the Clinical Proteomic Tumor Analysis Consortium (CPTAC), The Cancer Genome Atlas (TCGA) and the Camelyon 16 (CM16) were used.

| (a) CPTAC |  |
| --- | --- |
| Description | Value |
| Number of patients | 333 |
| Number of WSIs | 1336 |
| Number of WSIs per class<sup>1</sup> | 657, 679 |
| Number of patches<sup>2</sup> | 5885.0, [180-66011] |
| Age<sup>3</sup> | 0, 4, 22, 76, 129, 91, 5, 6 |
| Gender<sup>4</sup> | 229, 104 |
| Race<sup>5</sup> | 92, 6, 1, 1, 230 |

| (b) TCGA-NSCLC |  |
| --- | --- |
| Description | Value |
| Number of patients | 919 |
| Number of WSIs | 1053 |
| Number of WSIs per class<sup>1</sup> | 541, 512 |
| Number of patches<sup>2</sup> | 11604.5, [162-46625] |
| Age<sup>3</sup> | 0, 3, 42, 163, 325, 313, 54, 19 |
| Gender<sup>4</sup> | 558, 361 |
| Race<sup>5</sup> | 667, 80, 16, 1, 155 |

| (c) CM16 |  |
| --- | --- |
| Description | Value |
| Number of patients | 400 |
| Number of WSIs | 400 |
| Number of WSIs per class<sup>6</sup> | 240, 160 |
| Number of patches<sup>2</sup> | 10372.0, [142-55852] |
| Age<sup>3</sup> | N/A |
| Gender<sup>4</sup> | N/A |
| Race<sup>5</sup> | N/A |

<sup>1</sup> LUAD, LSCC
<sup>2</sup> Median, Range <sup>3</sup> Binned: 20-29, 30-39, 40-49, 50-59, 60-69, 70-79, ≥80, Unknown <sup>4</sup> Males, Females
<sup>5</sup> White, Black or African-American, Asian, American Indian or Alaskan Native, Other/unknown
<sup>6</sup> Normal, nodal metastases

#### CAMELYON16

[37] focuses on identifying breast cancer metastasis in WSIs of sentinel axillary lymph nodes, consisting of 400 WSIs with 270 for training and 130 for testing, noting one duplicate in the test set. The training set comprises 159 normal and 111 tumor-containing slides. The dataset’s challenge, as highlighted by prior studies [4], [19], lies in its small regions of metastatic cancer cells amidst extensive tissue areas, necessitating precise tumor cell detection. We divided the training set into new training and validation sets in a 4 : 1 ratio and tested all methods on the official test set, conducting a with versus without nodal metastases classification.

#### TCGA-NSCLC

[38] offers 1053 diagnostic slides, including 541 LUAD (lung adenocarcinoma) slides from 443 cases and 512 LUSC (lung squamous cell carcinoma) slides from 476 cases, representing two non-small cell lung cancer subtypes within the dataset. Tumor regions in TCGA-NSCLC are notably larger than in cancer cell clusters in CAMELYON16. We partitioned this dataset into training, validation, and testing subsets in a 3 : 1 : 1 ratio, using it for binary classification between LUAD and LUSC.

#### CPTAC

[39], comprising 1, 336 slides with 657 LUAD and 679 LUSC slides, presents tumor regions akin in size to TCGA-NSCLC. It served as an external dataset to evaluate the generalizability of models trained on the TCGA-NSCLC training dataset.

Post-processing yielded an average of 11, 556 patches per slide for CAMELYON16, 12, 261 for TCGA, and 7, 526 for CPTAC. For brevity, we refer to CAMELYON16 as CM16 and TCGA-NSCLC as TCGA throughout the remainder of this document.

### B. Implementation

#### Performance metrics

Our evaluation framework utilized the area under the receiver operating characteristic curve (AUC) and accuracy scores as primary metrics. The AUC score provides a comprehensive measure of the model’s capability to distinguish among various classes. We employed 5-fold cross-validation to ensure the reliability and generalizability of our results.

#### Training settings

We benchmarked our model against state-of-the-art (SOTA) multiple instance learning (MIL) approaches, including attention-based models like ABMIL [2], CLAM-SB/CLAM-MB with a single or multiple attentions branch [3], DTFD-MIL featuring a double-tier framework [4], as well as graph-based Patch-GCN [32], GTP [33], and the transformer-based TransMIL [19]. Whenever possible, official implementations were utilized to ensure fidelity in reproducing existing results. All models were evaluated using a consistent image resolution of 20X, in line with recommendations from the original publications.

#### Hyperparameters and training details

We opted for cross-entropy loss and employed the Lookahead optimizer [40], setting the learning rate to 2 ×10^−4^ and the weight decay to 1 × 10^−5^. The batch size was set to 1. Consistent with SOTA methods, we applied a fully connected layer to compress each feature embedding from 1024 to 512 dimensions, resulting in a hidden dimension (*D*) of 512. Our model incorporated 2 APFF blocks and was developed using PyTorch. Training was conducted on a single NVIDIA GTX 2080Ti GPU.

## V. Results and discussion

### A. Comparison with state-of-the-art methods

Table II details the performance comparison of our method against current SOTA models, showing our approach leads in accuracy across all datasets and achieves top AUC scores. The challenge presented by the CAMELYON16 dataset is notable due to its minimal regions of cancer cells (average total cancer cell area per slide *<* 10%), impacting the effectiveness of detection models. Our method, however, demonstrates significant improvements over existing models in both accuracy (by at least 1.9% point) and AUC scores (by 3.0% point). This suggests FourierMIL is capable of detecting sparse tumor signals by identifying correlations between tokens in the frequency domain. In the TCGA-NSCLC dataset, our method surpasses other methods, indicating at least a 1.6% point increase in accuracy and a 0.5% point rise in AUC scores. For the CPTAC dataset, our model outperforms competitors by at least 1.3% point in accuracy and 0.5% point in AUC scores. The effectiveness of our approach can be attributed to the APFF module’s ability to capture global dependencies and complex patterns in WSIs through efficient token mixing in the frequency domain.

**TABLE II:**
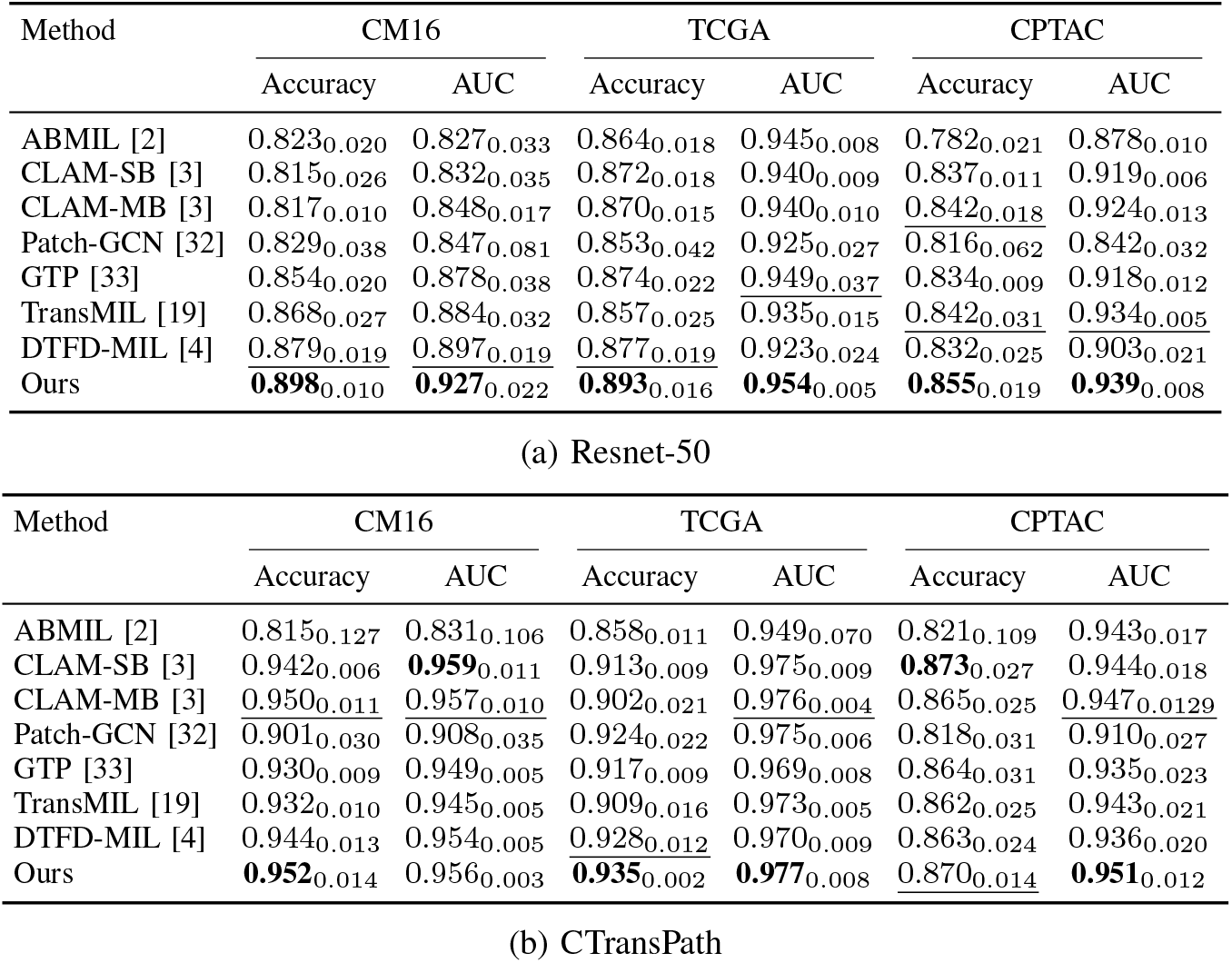
Classification results on Camelyon16, TCGA-NSCLC, CPTAC using (a) Resnet-50 and (b) CTransPath as the feature extractors. The best scores are in bold, second best results are underlined. The subscript in each cell is the standard derivation.

### B Ablation studies

#### APFF in token mixing and channel mixing layers

The impact of our APFF token mixing was assessed by contrasting the FourierMIL model with and without APFF token mixing layers. The results, presented in Table III, reveal that including APFF token mixing significantly enhances both accuracy and AUC scores on the Camelyon16 dataset by 27.8% and 43.4% point, respectively, compared to the model lacking APFF, which inaccurately classified all test samples as not have nodal metastases. Given that the latter model relies solely on channel mixing layers, we further explored their contribution by evaluating a model devoid of channel mixing layers, referred to as the “w/o CM” model. This investigation showed improvements of 9.6% point in accuracy and 9.0% point in AUC scores due to channel mixing layers. These findings underscore the utility of both APFF token mixing and channel mixing layers in our MLP-based FourierMIL for analyzing WSIs. Additionally, we compared filtering operations in both the original and frequency domains to elucidate the advantages of operating in the frequency domain. By eliminating all real-time Fourier and inverse real-time Fourier transformations in our FourierMIL, resulting in the “w/o FT” model, we observed a performance decline to predicting all test samples as normal, similar to the “w/o APFF” model. This indicates that APFF mixing’s implementation in the frequency domain is crucial for effective global token mixing.

**TABLE III:** Comparison of different modules in FourierMIL.

| Method | Accuracy | AUC |
| --- | --- | --- |
| w/o CM | 0.802<sub>0.034</sub> | 0.837<sub>0.025</sub> |
| w/o APFF | 0.620<sub>0.000</sub> | 0.493<sub>0.014</sub> |
| w/o FT | 0.620<sub>0.000</sub> | 0.505<sub>0.014</sub> |
| FourierMIL | 0.898<sub>0.010</sub> | 0.927<sub>0.022</sub> |

#### Effectiveness of all-pass filtering

We investigated the impact of employing low-pass (LP), high-pass (HP), and all-pass (AP) filters in our model’s frequency domain, as depicted in Table IV. Performance on the CM16 and TCGA datasets with LP and HP filters highlight the importance of both low-frequency and high-frequency components in classification tasks. Low-frequency elements within tokens help capture long-term trends and broad patterns, while high-frequency elements are crucial for identifying rapid changes, detailed nuances, and transient features within the tokens. Specifically, in the CM16 dataset, where tumor regions are a small fraction of the WSIs, HP filtering was more effective in distinguishing between lymph node tissue and cancer cell clusters, improving accuracy by 2.2% point for CM16. In contrast, the TCGA dataset, characterized by larger tumor regions, showed similar accuracy figures across filters. Using AP filtering allowed our model to utilize both low and high-frequency information, resulting in accuracy increases of at least 1.1% and 0.6% point for CM16 and TCGA, respectively.

**TABLE IV:** Ablation studies on all-pass (AP), low-pass (LP), and high-pass (HP). *λ* is set to 0.02 for LP and HP.

| Method | CM16 | TCGA |
| --- | --- | --- |
| LP | 0.865<sub>0.013</sub> | 0.887<sub>0.013</sub> |
| HP | 0.887<sub>0.022</sub> | 0.885<sub>0.014</sub> |
| AP | 0.898<sub>0.010</sub> | 0.893<sub>0.016</sub> |

#### Effect of adaptive token padding

The results in Table V illustrate the effects of employing various token padding (TP) strategies. Compared to the model without TP, introducing TP improved model performance, for instance, yielding an increase of 3.8% point in accuracy for CM16 and 1.3% point for TCGA. This finding supports the hypothesis that TP can mitigate the adverse effects of Fourier transform on image token mixing. Given the variability in token numbers across samples, we opted for a fixed ratio of token numbers to determine the padding amount for each sample, comparing these results with adaptive token padding (ATP). According to Table V, ATP outperformed the 10% *TP* strategy by a 1.7% point margin in accuracy for CM16, though it showed a slight reduction of 0.2% point compared to 10% *TP*. Token padding involves concatenating two tensors along a specified dimension without changing the tensor elements, prompting an evaluation of memory usage for different padding strategies. Findings indicated that ATP consumed less memory than 5% *TP*, suggesting that the average padding was below 5%, yet ATP provided better or comparable performance with reduced memory usage. These observations confirm ATP as a resource-efficient and effective approach to token padding.

**TABLE V:** Benefit of ATP. Average accuracy scores and standard deviations (subscript) are reported.

| Method | MEM(gb) | CM16 | TCGA |
| --- | --- | --- | --- |
| w/o TP | 0.309 | 0.860<sub>0.020</sub> | 0.880<sub>0.018</sub> |
| 5% TP | 0.332 | 0.867<sub>0.025</sub> | 0.891<sub>0.022</sub> |
| 10% TP | 0.351 | 0.881<sub>0.015</sub> | 0.895<sub>0.019</sub> |
| 20% TP | 0.377 | 0.878<sub>0.019</sub> | 0.884<sub>0.020</sub> |
| ATP | 0.324 | 0.898<sub>0.010</sub> | 0.893<sub>0.016</sub> |

#### Comparison with different SOTA as token mixing in FourierMIL

We evaluated the performance of our APFF module against that of other advanced methods, such as Nyströmformer and AFNO, focusing on their impact on token mixing. While most MLP-based methods are tailored for 2D input features and thus not directly comparable due to their reliance on 2D convolution operations, we assessed transformer-based approaches that employ linear attention mechanisms due to the high token count per sample. However, memory constraints limited the feasibility of using many linear attention transformers. Consequently, we selected Nyströmformer, known for its transformer-based architecture, and AFNO, which applies frequency filtering in one-dimensional scenarios, for direct comparison. Our comparative analysis involved substituting our APFF token mixing layer with analogous operations from these selected methods. The comparison utilized a baseline model, denoted as *Base*., representing our FourierMIL without the APFF layer. According to the results in Table VI, token mixing operations in the real domain, exemplified by *Base*. + Nyström attention, did not perform as well as our frequency domain approach. While AFNO also operates in the frequency domain and employs adaptive token filtering, it overlooks crucial information across various frequencies, particularly in the high-frequency range. Our analysis showed that our FourierMIL outperforms the *Base*. + AFNO configuration in both accuracy and AUC scores by 3.1% and 1.0% point, respectively. This outcome highlights the effectiveness of APFF’s all-pass function in capturing comprehensive frequency domain information. It is important to acknowledge a limitation: the Fourier transform operates at a computational complexity of O(n log n), while Nyströmformer achieves the more favorable O(n) time complexity.

**TABLE VI:** Comparison of SOTA models as token mixing in FourierMIL.

| Method | Accuracy | AUC |
| --- | --- | --- |
| *Base.*+Nyström [20] | 0.870<sub>0.031</sub> | 0.875<sub>0.045</sub> |
| *Base.*+AFNO [26] | 0.867<sub>0.028</sub> | 0.917<sub>0.009</sub> |
| *Base.*+APFF | 0.898<sub>0.010</sub> | 0.927<sub>0.022</sub> |

### C. Model interpretability

Model alignment with expert annotations in the relevant domain is fundamental to its effectiveness. We highlighted the interpretability of our approach by comparing our model’s outputs with reference annotations provided by expert pathologists, as seen in the CAMELYON16 dataset WSIs. Figure 2 showcases WSIs of alongside heatmaps generated from attention weights, enabling a direct comparison between our model and the CLAM model in identifying and interpreting regions of interest (ROI) and crucial morphological features for diagnosis. Figure 2 (a-c) demonstrates our model’s precision in identifying salient features within WSIs, closely matching the pathologist’s annotations. Conversely, Fig. 2(d-f) highlights a case with tumor presence less than 1% of the tissue region, where our model accurately delineates the tumor area with high precision and a minimal false positive rate. This comparative analysis highlights our model’s alignment with pathologist assessments and improved performance over CLAM in minimizing false positives. However, it also reveals a limitation: while our method demonstrates high precision, it exhibits low recall, indicating room for improvement in detecting all relevant cases.

**Fig. 2:**
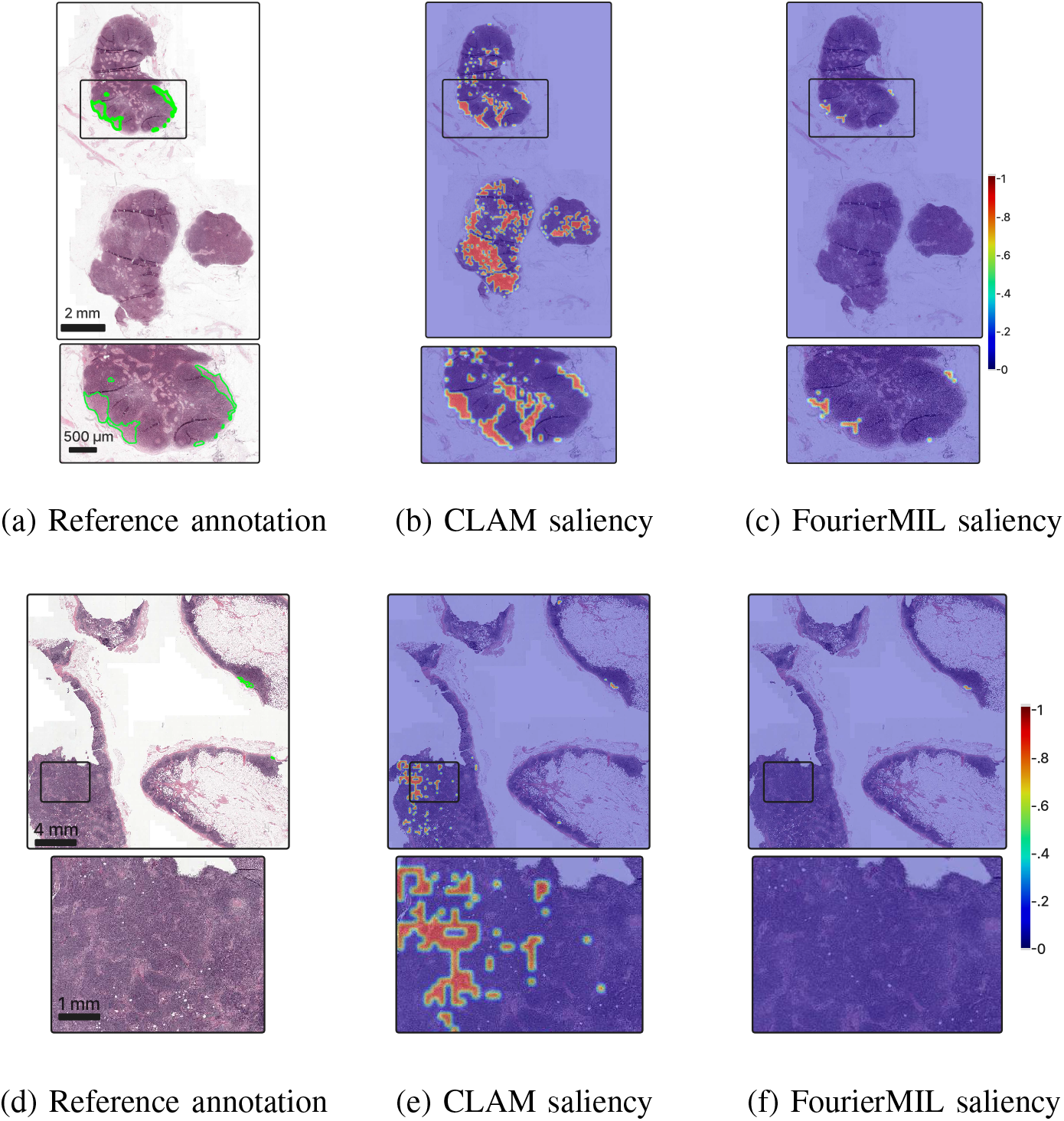
Visualization of nodal metastases detection. Panels (a-c) compare the reference annotations with CLAM and FourierMIL saliency heatmaps between two cases, highlighting annotations in green. Panels (d-f) depict a challenging case with a small ROI (< 1%), showcasing the model’s precision and the low rate of false positives in detecting subtle cancerous regions. The heatmap color intensities reflect each model’s attention to the detected areas (Created using QuPath [41]).

## VI. Conclusion

Our study introduces a novel approach to histopathological image analysis through the implementation of an all-pass frequency filtering (APFF) module, designed to enhance token mixing efficiency within WSIs. By evaluating the module’s performance across various datasets and against existing methods, we observed a consistent improvement in accuracy and area under the curve metrics. This improvement is particularly notable in challenging datasets characterized by sparse tumor regions, where our method’s ability to capture global dependencies and subtle details in the frequency domain proves advantageous. The comparative analysis with state-of-the-art methods, including Nyströmformer and AFNO, further underscores the unique benefits of applying APFF in the frequency domain for histopathological analysis. Our findings suggest that FourierMIL could play a critical role in advancing the accuracy of tumor detection and classification, thereby supporting more precise diagnostics.

## References

[1] K. He, X. Zhang, S. Ren, and J. Sun, “Deep Residual Learning for Image Recognition,” 2016 IEEE Conference on Computer Vision and Pattern Recognition (CVPR), pp. 770–778, 2015.

[2] M. Ilse, J. Tomczak, and M. Welling, “Attention-based Deep Multiple Instance Learning,” in Proceedings of the 35th International Conference on Machine Learning, J. Dy and A. Krause, Eds. 10–15 Jul 2018, vol. 80 of Proceedings of Machine Learning Research, pp. 2127–2136, PMLR.

[3] M. Y. Lu et al., “Data-efficient and weakly supervised computational pathology on whole-slide images,” Nature Biomedical Engineering, vol. 5, no. 6, pp. 555–570, 2021.

[4] H. Zhang, Y. Meng, Y. Zhao, Y. Qiao, X. Yang, S. E. Coupland, and Y. Zheng, “DTFD-MIL: Double-tier feature distillation multiple instance learning for histopathology whole slide image classification,” in 2022 IEEE/CVF Conference on Computer Vision and Pattern Recognition (CVPR), 2022, pp. 18780–18790.

[5] A. Krizhevsky, I. Sutskever, and G. E. Hinton, “ImageNet Classification with Deep Convolutional Neural Networks,” in Advances in Neural Information Processing Systems, F. Pereira, C. Burges, L. Bottou, and K. Weinberger, Eds. 2012, vol. 25, Curran Associates, Inc.

[6] K. Simonyan and A. Zisserman, “Very Deep Convolutional Networks for Large-Scale Image Recognition,” in 3rd International Conference on Learning Representations, ICLR 2015, San Diego, CA, USA, May 7-9, 2015, Conference Track Proceedings, Y. Bengio and Y. LeCun, Eds., 2015.

[7] J. Saltz et al., “Spatial Organization and Molecular Correlation of Tumor-Infiltrating Lymphocytes Using Deep Learning on Pathology Images,” Cell Reports, vol. 23, no. 1, pp. 181–193.e7, 2018.

[8] A. Vaswani, N. Shazeer, N. Parmar, J. Uszkoreit, L. Jones, A. N. Gomez, L. Kaiser, and I. Polosukhin, “Attention Is All You Need,” CoRR, vol. abs/1706.03762, 2017.

[9] A. Dosovitskiy, L. Beyer, A. Kolesnikov, D. Weissenborn, X. Zhai, T. Unterthiner, M. Dehghani, M. Minderer, G. Heigold, S. Gelly, J. Uszkoreit, and N. Houlsby, “An Image is Worth 16×16 Words: Transformers for Image Recognition at Scale,” in International Conference on Learning Representations, 2021.

[10] I. O. Tolstikhin, N. Houlsby, A. Kolesnikov, L. Beyer, X. Zhai, T. Unterthiner, J. Yung, D. Keysers, J. Uszkoreit, M. Lucic, and A. Dosovitskiy, “MLP-Mixer: An all-MLP Architecture for Vision,” in Neural Information Processing Systems, 2021.

[11] G. Campanella, M. G. Hanna, L. Geneslaw, A. Miraflor, V. Werneck Krauss Silva, K. J. Busam, E. Brogi, V. E. Reuter, D. S. Klimstra, and T. J. Fuchs, “Clinical-grade computational pathology using weakly supervised deep learning on whole slide images,” Nature Medicine, vol. 25, no. 8, pp. 1301–1309, 2019.

[12] G. Xu, Z. Song, Z. Sun, C. Ku, Z. Yang, C. Liu, S. Wang, J. Ma, and W. Xu, “CAMEL: A weakly supervised learning framework for histopathology image segmentation,” 2019 IEEE/CVF International Conference on Computer Vision (ICCV), pp. 10681–10690, 2019.

[13] N. Hashimoto, D. Fukushima, R. Koga, Y. Takagi, K. Ko, K. Kohno, M. Nakaguro, S. Nakamura, H. Hontani, and I. Takeuchi, “Multi-scale Domain-adversarial Multiple-instance CNN for Cancer Subtype Classification with Unannotated Histopathological Images,” in 2020 IEEE/CVF Conference on Computer Vision and Pattern Recognition (CVPR), Los Alamitos, CA, USA, jun 2020, pp. 3851–3860, IEEE Computer Society.

[14] B. Li, Y. Li, and K. W. Eliceiri, “Dual-stream multiple instance learning network for whole slide image classification with self-supervised contrastive learning,” in Proceedings of the IEEE/CVF Conference on Computer Vision and Pattern Recognition, 2021, pp. 14318–14328.

[15] Y. Sharma, A. Shrivastava, L. Ehsan, C. A. Moskaluk, S. Syed, and D. Brown, “Cluster-to-Conquer: A Framework for End-to-End Multi-Instance Learning for Whole Slide Image Classification,” in Medical Imaging with Deep Learning, 2021.

[16] X. Wang et al., “Weakly Supervised Deep Learning for Whole Slide Lung Cancer Image Analysis,” IEEE Transactions on Cybernetics, vol. 50, no. 9, pp. 3950–3962, 2020.

[17] J. Deng, W. Dong, R. Socher, L.-J. Li, K. Li, and F.-F. Li, “ImageNet: a Large-Scale Hierarchical Image Database,” 06 2009, pp. 248–255.

[18] S. Kalra, M. Adnan, G. Taylor, and H. R. Tizhoosh, “Learning Permutation Invariant Representations Using Memory Networks,” in European Conference on Computer Vision. 2020, pp. 677–693, Springer.

[19] Z. Shao, H. Bian, Y. Chen, Y. Wang, J. Zhang, X. Ji, and Y. Zhang, “TransMIL: Transformer based Correlated Multiple Instance Learning for Whole Slide Image Classification,” in Advances in Neural Information Processing Systems, M. Ranzato, A. Beygelzimer, Y. Dauphin, P. Liang, and J. W. Vaughan, Eds. 2021, vol. 34, pp. 2136–2147, Curran Associates, Inc.

[20] Y. Xiong, Z. Zeng, R. Chakraborty, M. Tan, G. Fung, Y. Li, and V. Singh, “Nyströmformer: A Nyström-based Algorithm for Approximating Self-Attention,” Proceedings of the AAAI Conference on Artificial Intelligence, vol. 35, no. 16, pp. 14138–14148, May 2021.

[21] S. Chen, E. Xie, C. Ge, R. Chen, D. Liang, and P. Luo, “CycleMLP: A MLP-like architecture for dense visual predictions,” IEEE Trans. Pattern Anal. Mach. Intell., vol. 45, no. 12, pp. 14284–14300, 2023.

[22] D. Lian, Z. Yu, X. Sun, and S. Gao, “AS-MLP: An axial shifted MLP architecture for vision,” in The Tenth International Conference on Learning Representations, ICLR 2022, Virtual Event, April 25-29, 2022. 2022, OpenReview.net.

[23] T. Yu, X. Li, Y. Cai, M. Sun, and P. Li, “S2-MLP: Spatial-Shift MLP Architecture for Vision,” in Proceedings - 2022 IEEE/CVF Winter Conference on Applications of Computer Vision, WACV 2022, United States, 2022, Proceedings - 2022 IEEE/CVF Winter Conference on Applications of Computer Vision, WACV 2022, pp. 3615–3624, Institute of Electrical and Electronics Engineers Inc.

[24] J. Lee-Thorp, J. Ainslie, I. Eckstein, and S. Ontanon, “FNet: Mixing Tokens with Fourier Transforms,” in Proceedings of the 2022 Conference of the North American Chapter of the Association for Computational Linguistics: Human Language Technologies, M. Carpuat, M.-C. de Marneffe, and I.V. Meza Ruiz, Eds., Seattle, United States, July 2022, pp. 4296–4313, Association for Computational Linguistics.

[25] Y. Rao, W. Zhao, Z. Zhu, J. Lu, and J. Zhou, “Global filter networks for image classification,” in Advances in Neural Information Processing Systems (NeurIPS), 2021.

[26] J. Guibas, M. Mardani, Z. Li, A. Tao, A. Anandkumar, and B. Catanzaro, “Efficient Token Mixing for Transformers via Adaptive Fourier Neural Operators,” in International Conference on Learning Representations, 2021.

[27] Y. Zhao, F. Yang, Y. Fang, H. Liu, N. Zhou, J. Zhang, J. Sun, S. Yang, B. Menze, X. Fan, and J. Yao, “Predicting Lymph Node Metastasis Using Histopathological Images Based on Multiple Instance Learning With Deep Graph Convolution,” in 2020 IEEE/CVF Conference on Computer Vision and Pattern Recognition (CVPR), 2020, pp. 4836–4845.

[28] W. Lu, S. Graham, M. Bilal, N. Rajpoot, and F. Minhas, “Capturing Cellular Topology in Multi-Gigapixel Pathology Images,” in 2020 IEEE/CVF Conference on Computer Vision and Pattern Recognition Workshops (CVPRW), 2020, pp. 1049–1058.

[29] Y. Zhou, S. Graham, N. Alemi Koohbanani, M. Shaban, P.-A. Heng, and N. Rajpoot, “CGC-Net: Cell Graph Convolutional Network for Grading of Colorectal Cancer Histology Images,” in 2019 IEEE/CVF International Conference on Computer Vision Workshop (ICCVW), 2019, pp. 388–398.

[30] R. J. Chen et al., “Pathomic Fusion: An Integrated Framework for Fusing Histopathology and Genomic Features for Cancer Diagnosis and Prognosis,” IEEE Transactions on Medical Imaging, vol. 41, no. 4, pp. 757–770, 2022.

[31] R. Li, J. Yao, X. Zhu, Y. Li, and J. Huang, “Graph CNN for Survival Analysis on Whole Slide Pathological Images,” in Medical Image Computing and Computer Assisted Intervention – MICCAI 2018, A. F. Frangi, J. A. Schnabel, C. Davatzikos, C. Alberola-López, and G. Fichtinger, Eds., Cham, 2018, pp. 174–182, Springer International Publishing.

[32] R. J. Chen, M. Y. Lu, M. Shaban, C. Chen, T. Y. Chen, D. F. Williamson, and F. Mahmood, “Whole Slide Images are 2D Point Clouds: Context-Aware Survival Prediction using Patch-based Graph Convolutional Networks,” in Medical Image Computing and Computer Assisted Intervention – MICCAI 2021, pp. 339–349. Springer International Publishing, 2021.

[33] Y. Zheng, R. H. Gindra, E. J. Green, E. J. Burks, M. Betke, J. E. Beane, and V. B. Kolachalama, “A Graph-Transformer for Whole Slide Image Classification,” IEEE Transactions on Medical Imaging, vol. 41, no. 11, pp. 3003–3015, 2022.

[34] X. Wang, S. Yang, J. Zhang, M. Wang, J. Zhang, W. Yang, J. Huang, and X. Han, “Transformer-based unsupervised contrastive learning for histopathological image classification,” Medical Image Analysis, vol. 81, pp. 102559, 2022.

[35] S. S. Soliman and M. D. Srinath, Continuous and discrete signals and systems, SAO/NASA, 1990.

[36] D. Hendrycks and K. Gimpel, “Bridging nonlinearities and stochastic regularizers with Gaussian error linear units,” 2017.

[37] B. Ehteshami Bejnordi et al., “Diagnostic Assessment of Deep Learning Algorithms for Detection of Lymph Node Metastases in Women With Breast Cancer,” JAMA, vol. 318, no. 22, pp. 2199–2210, 2017.

[38] TCGA Research Network, “The Cancer Genome Atlas Program,” Available: https://portal.gdc.cancer.gov/.

[39] N. J. Edwards, M. Oberti, R. R. Thangudu, S. Cai, P. B. McGarvey, S. Jacob, S. Madhavan, and K. A. Ketchum, “The CPTAC Data Portal: A Resource for Cancer Proteomics Research,” Journal of Proteome Research, vol. 14, no. 6, pp. 2707–2713, 2015.

[40] M. R. Zhang, J. Lucas, G. E. Hinton, and J. Ba, “Lookahead optimizer: k steps forward, 1 step back,” in Advances in Neural Information Processing Systems, 2019, pp. 9597–9608.

[41] F. J. e. a. Bankhead P., Loughrey M.B., “Qupath: Open source software for digital pathology image analysis.,” in Scientific Reports, 2017.

